# A 3D-printed handheld device for quick citrus tissue lysis and nucleic acid extraction

**DOI:** 10.1101/2024.08.26.609775

**Authors:** Chia-Wei Liu, Brent Kalish, Sohrab Bodaghi, Georgios Vidalakis, Hideaki Tsutsui

**Affiliations:** Department of Mechanical Engineering, University of California, Riverside, CA, 92521, USA; Department of Microbiology and Plant Pathology; University of California, Riverside, CA, 92521, USA; Department of Bioengineering; University of California, Riverside, CA, 92521, USA

**Keywords:** nucleic acid extraction, sample preparation, 3D-printed handheld device, paper disk, Citrus tristeza virus, Spiroplasma citri

## Abstract

A 3D-printed handheld device has been developed for rapid and efficient sample preparation from citrus leaves, aimed at streamlining protocols traditionally reliant on mortar and pestle. With its high-speed motor, knurled lysis chamber for rapid sample lysis, and quick nucleic acid extraction using paper disks, this device can yield ready-to-use extracts in just 12 minutes, significantly reducing the time required for sample preparation. The device was optimized for maximum sample lysis by evaluating operation voltages and chamber features. The results showed that the lysis chamber with internal knurling and the motor operated at 7.5 V was sufficient for effective sample lysis in 1 minute, achieving total RNA concentrations up to 87.6% of those obtained with a mortar and pestle. Furthermore, concerns regarding heat generation and resin release during the lysis process were found to not impact sample quality. To further facilitate in-field diagnosis, the capability of in-device sample preparation was verified with citrus sources infected with *citrus tristeza virus* and *Spiroplasma citri* in qPCR-based assays, where low assay variations were demonstrated (< 3.8%). Overall, the in-device sample preparation integrated with the paper disks showed good reliability and compatibility across different pathogens for downstream analysis. An eco-friendly sterilization protocol using household bleach and vitamin C solution was also developed to safely reuse the device for in-field deployment.

## 1. Introduction

Plant epidemics have long been a problem affecting crop yields and the global economy, and it gets even more challenging for disease management as time passes due to the emergence of novel pathogenic variants and their disease synergism (Tatineni and Hein 2023). Specifically, these destructive plant diseases caused by different pathogens and pests lead to global crop yield losses of up to 40% for maize, potato, rice, soybean, and wheat, amounting to an annual financial loss of $220 billion (He and Krainer 2020; Singh et al., 2023). According to the UN’s 2024 Global Report on Food Crises, almost 282 million people across 59 countries/territories are now experiencing acute food insecurity, which has been increasing yearly since 2019 (World Food Programme 2024). Among the drivers causing the food crisis, crop loss due to extreme climate and plant pathogens will further exacerbate affected regions’ environmental and socio-economic conditions and disproportionately affect food-insecure populations (Ristaino et al., 2021; Singh et al., 2023). Take citrus, one of the most common cash crops in the world, as an example. As a mainstay in agricultural industries of many countries, the US, for example, has produced 4.9 million tons in the season 2022-2023, accounting for 2.58 billion USD (United States Department of Agriculture 2023). However, huanglongbing (HLB), a destructive citrus disease with rapid disease progression, has already severely reduced citrus yields due to the early abortion of fruits from affected branches (Gottwald 2010). Especially in Florida, the first state in the US to report HLB, citrus yield has been reduced by 80%, with tens of billions USD of economic losses in the past two decades (Alvarez et al., 2016; Graham et al., 2024). Therefore, being able to detect a disease in its early stages is of great importance for growers to prevent disease outbreaks and accurately adopt control strategies.

The current strategy for disease management relies on sensitive and high-throughput diagnostic methods, such as high-throughput sequencing (HTS),, third-generation sequencing (TGS), nanopore sequencing, and quantitative polymerase chain reaction (qPCR), to precisely identify diseased trees (Aragona et al., 2022; Chalupowicz et al., 2019; Sharma and Sharma 2016). It is followed by risk-reducing measures, including removal of the diseased plants and subsequent vector management (Tatineni and Hein 2023). However, this practice is insufficient for disease control because these sensing techniques require high-quality samples and are always tied to lab facilities, which are usually costly, time-consuming, and require trained personnel (Paul et al., 2020b). In particular, tedious specimen collection, packaging, and submission in the field by growers prior to sample preparation in labs is usually required (California Department of Food & Agriculture). At the lab, the procedures include multiple steps, such as manual tissue homogenization and nucleic acid extraction, followed by downstream assays like qPCR and sequencing (Dang et al., 2022). The entire process can take from a few days to several weeks, depending on the schedule and the specimen amount. Although this standard practice provides reliable results due to high-quality nucleic acids, the long turnaround times and labor intensiveness make it inefficient for disease control (Liu and Tsutsui 2023). The use of high-throughput instruments mitigates the problems by offering semi-automated and large-scale sample processing (Dang et al., 2022). Still, time-consuming sample pretreatment and high power requirements make them impractical for quick and on-site disease diagnosis (Dang et al., 2022). So far, some commercial products and lab studies, such as the QIAcard FTA PlantSaver and microneedle patch, have attempted to improve in-field sample preparation (Jia et al., 2021; Paul et al., 2020a; QIAGEN). However, the amount of nucleic acids these tools can extract is limited, which is unfavorable for early-stage detection. Serological tests, such as lateral flow immunoassays (e.g., AgriStrip, Pocket Diagnostics kits, etc.), could be a solution for in-field prescreening of plant diseases (Singh et al., 2021). However, follow-up testing of the positive samples with more specific techniques (e.g., qPCR, pathogen culture) in advanced labs would still be required (Miller et al., 2009; Singh et al., 2021).

To expedite the current practice and remove the need for manual grinding and costly lab instruments, a 3D-printed handheld device has been developed in this study to replace the current standard benchtop protocol of tissue homogenization and lysis via mortar and pestle (Fig. 1, left). This device, incorporating an efficient sample lysis unit and a quick nucleic acid extraction unit, has been optimized across multiple healthy and infected citrus species in terms of operation voltages and lysis chamber features. Specifically, a high-speed motor drives a 3D-printed blade to macerate citrus petioles and midveins in the lysis chamber and produce a crude lysate, followed by soaking paper disks in the lysate for quick nucleic acid extraction, enabling downstream molecular assays (Fig. 1, right). The handheld device protocol takes 10 minutes in total for the preparation of ready-to-use paper disks, where elution of total nucleic acids takes an additional 2 minutes for immediate on-site use. However, in order to meet the practical needs, prepared paper disks in this study were fully air-dried for 24 hours before elution as our anticipated use-case for this device is for dried, prepared paper disks to be shipped at room temperature from farmers to centralized labs for testing. As previously reported, an extreme case, such as the elution of air-dried paper disks stored at room temperature for a month, was still able to generate equivalent Cq values compared to fresh elution (Liu et al., 2024).

**Fig. 1.**
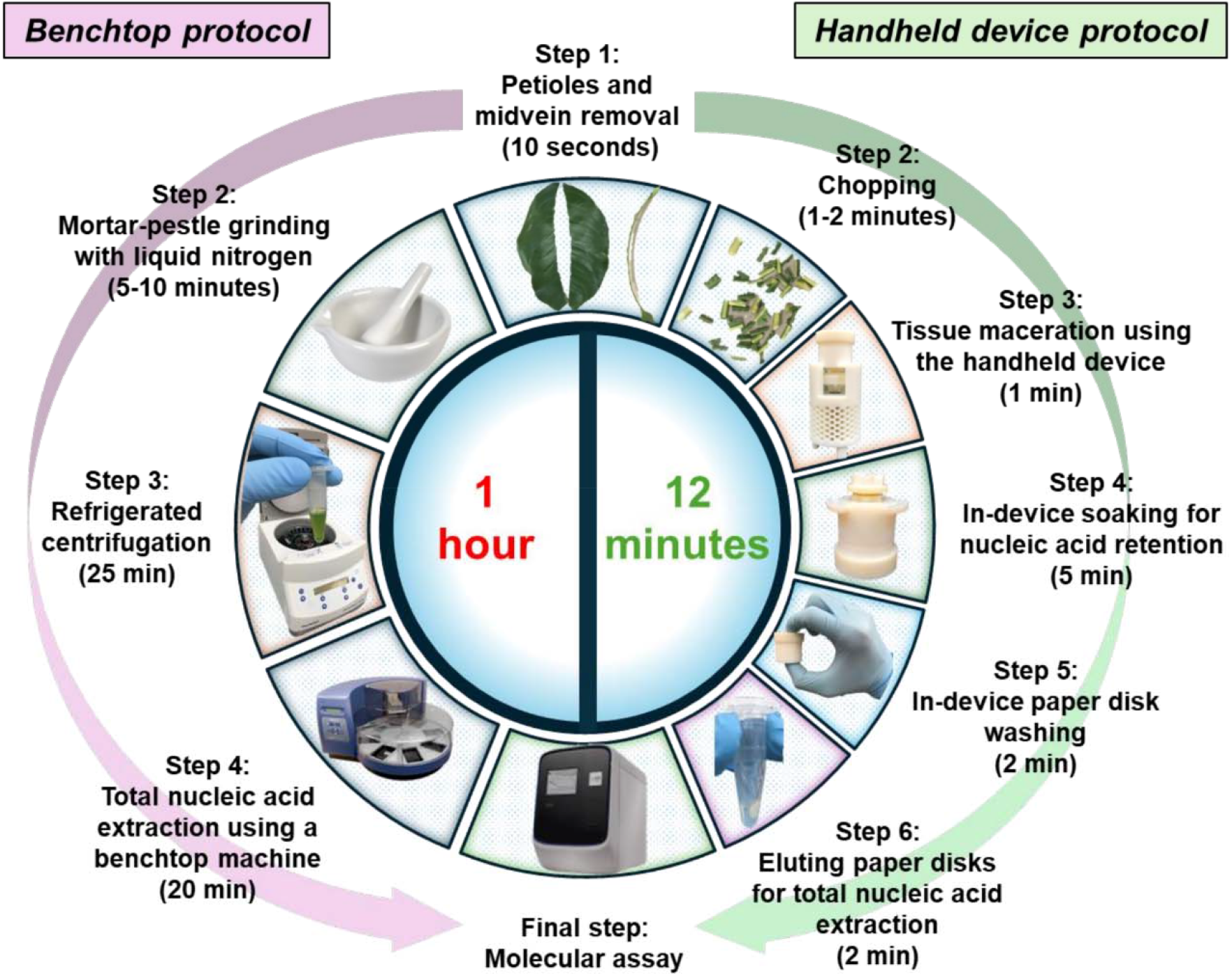
Stepwise comparison between a standard benchtop protocol (left) and the handheld device protocol proposed in this study (right), highlighting the key components for sample lysis and nucleic acid extraction.

This development aims to allow farmers and growers to prepare ready-to-use nucleic acids straight from plants of interest in a portable manner or retain them on dried paper disks for easier storage and shipping to labs. This will spare laboratory workers from processing large quantities of specimens and focus on harvesting nucleic acids from the dried paper disks for downstream analysis. The device reduces workloads and expedites the overall disease diagnosis process.

## 2. Material and methods

### 2.1 Materials and equipment

The handheld device was designed using SolidWorks 2022 (Dassault Systèmes SOLIDWORKS Corp., France) and printed in Veroclear-RGD810 and VeroWhitePlus-RGD835 with SUP705™ support in a Stratasys Objet30 3D printer (Stratasys, Ltd., Eden Prairie, MN). Total nucleic acid concentrations and purity were measured using a Qubit 4 fluorometer with RNA/dsDNA Broad Range (BR) assay kits (Cat. No. Q10210, Q32850, Life Technologies, Carlsbad, CA) and a NanoDrop 2000c spectrophotometer (Thermo Fisher Scientific, Waltham, MA), respectively. DNA/RNA extraction controls were done using a MagMAX Express-96 Magnetic Particle Processor with a MagMAX-96 Viral RNA isolation kit (Cat. No. AM1836) (Thermo Fisher Scientific, Waltham, MA). qPCR and RT-qPCR were performed in an Applied Biosystems QuantStudio 12K Flex Real-Time PCR System (Thermo Fisher Scientific, Waltham, MA). The electric motor used was an RS-380PH-4045 (Mabuchi Motor Company, Japan) and powered by a DIGI 35A DC power supply (Electro Industries, Monticello, MN). Motor speed was measured with an R7050 Compact Photo Tachometer (REED Instruments, Wilmington, NC). The handheld device’s shaft assembly was lubricated and sealed with silicone grease (Cat. No. 335148, Beckman Instruments, Palo Alto, CA). Temperatures inside the lysis chamber were measured using k-type thermocouples (Omega Engineering, Norwalk, CT) and an Arduino Uno Rev3 board (Arduino, Italy). ATR-FTIR spectra were obtained using Nicolet iS50 FTIR Advanced KBr Gold Spectrometer (Thermo Fisher Scientific, Waltham, MA).

### 2.2 Reagents and buffers

The lysis buffer used throughout the study was composed of 4 M guanidinium thiocyanate (GTC) (Sigma-Aldrich, St. Louis, MO), 0.2 M sodium acetate trihydrate (pH 5.0) (Thermo Fisher Scientific, Waltham, MA), 2 mM ethylenediaminetetraacetic acid (EDTA, Thermo Fisher Scientific, Waltham, MA), and 2.5% polyvinylpyrrolidone (PVP, Sigma-Aldrich, St. Louis, MO), where the final pH was adjusted to 5.0 with 1N hydrochloride (Liu et al., 2024). Paper disks (d = 0.25 in) used for quick nucleic acid extraction were cut from untreated Whatman Grade 1 CHR cellulose chromatography papers (Cat. No. 3001-861, Sigma-Aldrich, St. Louis, MO) using a hole puncher. Tris-HCl solution (1M, pH 7.5) (Cat. No. 15567-027, Invitrogen, Waltham, MA) was diluted to 10 mM to rinse the paper disks after soaking. Tris-EDTA (TE) 1X solution (pH 8.0) (Cat. No. BP2473100, Thermo Fisher Scientific, Waltham, MA) was used to resuspend nucleic acids after benchtop extraction. TE 1X solution with 100 mM sodium chloride (Thermo Fisher Scientific, Waltham, MA) added was used to elute paper disks. (Liu et al., 2024). Nuclease-free water (Milli-Q Biopak Polisher, Sigma-Aldrich, St. Louis, MO) was used to prepare MasterMix of qPCR-based assays. CloroxPro Clorox Germicidal Bleach (Clorox Healthcare, Oakland, CA) was used for disinfection of the handheld device, and sodium ascorbate (Eisen-Golden Store, Dublin, CA) was used to neutralize residual bleach in the wash step.

### 2.3 Overview of the handheld device

#### 2.3.1 Design and fabrication

All the parts comprising this handheld device were designed using SolidWorks 2022 and were fabricated with Veroclear-RGD810 and VeroWhitePlus-RGD835 build materials and SUP705™ support using Stratasys Objet30 3D printer. Sharp tools and brushes were used to remove most of the support, followed by a 5-minute sonication in 70% (v/v) ethanol and pure isopropanol sequentially before rinsing with water to remove organic solvents and particulates. The entire assembly with its exploded view of the three major parts (a paper-disk extraction unit, a lysis chamber, and a blade and clutch assembly) is depicted in Fig. 2a. The paper-disk extraction unit holds five untreated chromatography paper disks (d = 0.25 in.) cut using a hole puncher (Fig. 2b, i). The untreated Whatman grade 1 chromatography paper has previously been proven effective in retaining nucleic acids from crude lysates and releasing them back into an elution buffer (Liu et al., 2024). The specialized channel-hole design on the holder allows the wash buffer to flow easily back and forth and assists in removing tissue residues during the mild shake (Fig. 2b, i). The disk retainer holds the paper disks in place during soaking and washing steps (Fig. 2b, i). The lysis chamber is where the chopped tissues undergo 1-minute maceration with the lysis buffer. The knurling (Fig. 2b, ii) inside the chamber helps break down the tissues during the high-speed spinning. The safe lock around the chamber and the motor cap (Fig. 2b, iii) keeps the chamber fixed during operation. The blade and clutch assembly consist of a two-part clutch with the blades screwed into the upper clutch. The beveled surfaces of the clutch’s teeth help it fit into its drive counterpart more easily (Fig. 2b, iv). The beveled surfaces also are oriented against the direction of rotation, preventing slippage (Fig. 2b, iv). A cutaway drawing and a photo of the printed device are depicted in Fig. 2c.

**Fig. 2.**
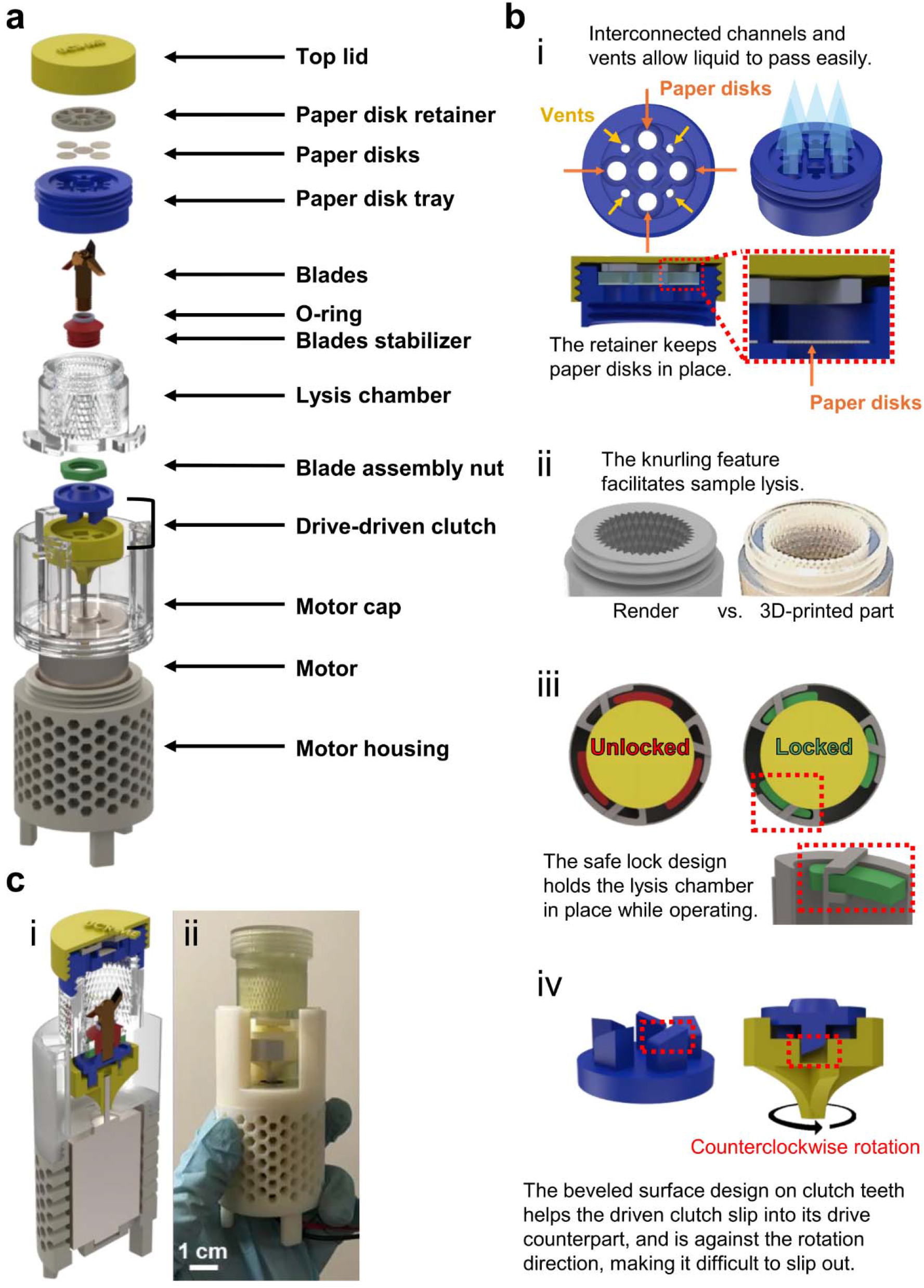
The complete design of the handheld device. (a) Exploded drawing of the handheld device with all parts labeled. (b) Selected major components shown to emphasize design concepts: i) Paper disk holder. The vent design allows fluids to flow back and forth and form agitation easily. A paper retainer keeps paper disks in place while fully interacting with fluids, ii) Knurled lysis chamber. The knurled inside of the chamber facilitates tissue maceration, iii) Lysis chamber safety lock. The easy safe-lock design holds the chamber in place while operating, iv) Clutch assembly. The beveled surface design on clutch teeth helps the clutch slip into its drive counterpart easily. The beveled surfaces are opposite the direction of rotation, preventing slippage. (c) The handheld device: i) cutaway drawing and ii) 3D-printed device.

#### 2.3.2 Operation procedure

The stepwise operation procedure of the handheld device is illustrated in Fig. S1. (a) Place 500 mg of chopped tissue (∼0.25 in. long) in the lysis chamber with 3000 µL guanidine lysis buffer, screw the paper disk unit and the top lid before switching on the power. (b) Power on for a 1-minute quick maceration and off. (c) Flip the device upside down for 5 minutes, soaking the paper disks in crude lysate, and then flip it upright. (d) Unscrew the lysis chamber and replace it with the wash chamber containing 1 mL of wash buffer. Mildly shake the assembly for 1 minute, making sure the wash buffer is sufficiently in contact with the paper disks. After 1 minute, discard the wash buffer and then repeat the process with 1mL of fresh wash buffer. (e) Move the paper disks to an aluminum foil sheet using clean tweezers and let them fully air dry (∼ 24 hours) before being ready to use. (f) Elute the paper disks to harvest total nucleic acids for downstream analysis.

### 2.4 Plant materials and sample collection

Healthy Sour orange (*Citrus aurantium* L.), healthy Washington navel (*Citrus sinensis* L. Osbeck), and healthy Limoneira (*Citrus limon* L. Burm.f.) were collected from Agriculture Operations (AgOps) at the University of California, Riverside (UCR). Healthy Sour orange was provided by the Citrus Clonal Protection Program (CCPP) at UCR. Diseased citrus sources, Pineapple sweet orange and Sweet orange (*Citrus sinensis* L. Osbeck), with single infection of *citrus tristeza virus* (CTV) and *Spiroplasma citri* (*S. citri*), respectively, were provided by the CCPP disease bank at the UCR (Table S1). The study used these sources to evaluate the developed handheld device and the corresponding protocol. All tested leaves were evenly collected from the east, west, south, and north sides of the tree to avoid uneven distribution of the pathogens and were gently wiped with wet paper napkins to remove dust on the leaves prior to processing.

### 2.5 Benchtop protocol

#### 2.5.1 Preparation of crude lysates

According to the protocol previously reported (Liu et al., 2024), phloem-rich petioles and midveins peeled off from the collected leaves were ground using mortar and pestle with the addition of liquid nitrogen until finely powdered. The ground tissues were then aliquoted to 100 mg, lysed with 600 µL of guanidine lysis buffer, and vortexed for 1 min, followed by 15 minutes of incubation at 4°C. The mixture was centrifuged at 14,000 × g at 4°C for 25 minutes, where 150 μL of the supernatant was collected for subsequent nucleic acid extraction.

#### 2.5.2 Nucleic acid extraction using benchtop instruments

Total nucleic acids in crude lysates were isolated using a pre-programmed MagMAX Express-96 Magnetic Particle Processor with a MagMAX-96 Viral RNA isolation kit. Each reaction contained 150 μL of supernatant of crude lysate, 22 μL of RNA binding bead mix (10 μL lysis/binding enhancer + 10 μL RNA binding beads + 2 μL carrier RNA), 139 μL of lysis/binding solution as recommended by the manufacturer and an extra 139 μL of 100% isopropanol as previously reported (Dang et al., 2022). The resulting nucleic acids were eluted in 100 μL of TE 1X solution and assessed using a NanoDrop 2000c spectrophotometer to confirm the quality. Nucleic acid concentrations (ng/μL) were determined using a Qubit 4 fluorometer with RNA/dsDNA Broad Range (BR) assay kits. These extracts were then aliquoted and stored at −80°C for later use.

### 2.6 Handheld device protocol

#### 2.6.1 Preparation of crude lysates

The proposed handheld device was used for lysate preparation throughout the study. Similarly, petioles and midveins were collected from leaves and chopped into a quarter-inch size with a clean razor blade to fit inside the lysis chamber of the device. The chopped tissues were weighed out to 500 mg each, then ground and lysed with 3000 µL of guanidine lysis buffer for 1 minute at the specified voltage. The lysate then underwent either a benchtop extraction, as described above, or a direct extraction using paper disks.

#### 2.6.2 Nucleic acid extraction using paper disks

The handheld device was assembled by inserting five paper disks at the beginning of the process. As previously reported (Liu et al., 2024), 5 minutes of paper-disk soaking is optimal for nucleic acid capture. Thus, the entire device was inverted to allow the paper disks to soak in the lysate for 5 minutes after homogenization. After soaking, the device was flipped back upright and the lysis chamber is replaced with a wash chamber containing 1 mL of 10 mM Tris-HCl solution (pH 7.5), followed by 1-minute of gentle shaking twice to clean the paper disks. Using sterilized tweezers, the five washed disks were transferred to an aluminum foil sheet for 24-hour air drying before eluting them in 100 µL of modified TE 1X solution for molecular assays. Nucleic acid concentrations (ng/μL) were determined using a Qubit 4 fluorometer with RNA/dsDNA Broad Range (BR) assay kits before storing at −80°C. These washed disks can also be air-dried and stored at room temperature for later use.

### 2.7 qPCR and RT-qPCR assays

Primers and probes used in the study for singleplex qPCR/RT-qPCR assays were adapted from previous reports (Osman et al., 2017; Osman et al., 2015; Shi et al., 2014). Primers and probes targeting the cytochrome oxidase (COX) gene in the citrus genome are routinely used as a reliable internal control to confirm nucleic acid integrity (Osman et al., 2017; Osman et al., 2015). The sequences of the primers, the probes, and the corresponding nucleotide positions of the targets of interest (i.e., CTV, *S. citri*, and COX) are available in Table S2. All the assays were conducted using an Applied Biosystems QuantStudio 12K Flex Real-Time PCR System. The specific kits and protocols used for singleplex qPCR/RT-qPCR assays are detailed in Table S3.

## 3. Results and discussion

### 3.1 Characterization of the handheld device

The high-speed motor driving the handheld device, RS-380PH-4045, has a voltage range of 4.5-9.6 V with a normal working voltage of 7.2 V. To determine the correlation between rotation speed and lysis efficiency for plant tissues, three voltages, 6, 7.5, and 9 V, equally distributed within the range, were selected for the series of assessments below. In addition, while buffer leaking from the interstice between the shaft and stabilizer was found to be an issue during practical operation, it was significantly reduced with the addition of silicone grease before each operation, as detailed in Fig. S2. Each operation’s rotation speed (in RPM) under different conditions was measured using a tachometer to identify any possible differences (Fig. S3).

#### 3.1.1 Evaluation of heat generation

Heat generation during the lysis process could inversely affect the quality of nucleic acids, especially RNAs, due to the digestion of more active enzymes (e.g., RNase) at a favorable temperature (Fan and Gulley 2001). In addition, elevated temperatures can also lead to the denaturation of RNA, unfolding secondary and tertiary structures. This can result in RNA hydrolysis and cause irreversible damage to RNA integrity (Becskei and Rahaman 2022). To evaluate heat generation during the 1-min lysis process under different operation voltages, the temperature was measured at the two spots marked in blue and red in Fig. 3a. using k-type thermocouples and an Arduino Uno Rev3 board. In this test, the lysis chamber was filled with 3 mL of ddH_2_O at room temperature (∼25 °C), replacing the guanidine lysis buffer to prevent the corroding, leading to inaccurate measurements (Pistorius and Li 2012). As shown in Fig. 3b-d, the temperature differences (ΔT) under 6V are 0.81±0.06 °C (top) and 1.08±0.19 °C (bottom), 0.96±0.24 °C (top) and 1.17±0.12 °C (bottom) under 7.5V, and 1.12±0.2 °C (top) 1.46±0.08 °C (bottom) under 9V. Although it is clear that ΔT increased with the operation voltage, the changes are very small. In addition, temperatures measured at the bottom were always slightly higher than those measured at the top, however, the differences were negligible (< 0.34 °C). It is inferred that heat is generated due to the high-speed spinning of the shaft in contact with the stabilizer at the bottom, where the temperature is transferred from. This assessment can be applied not only to guanidine lysis buffer but also to other water-based lysis buffers with higher specific heat, where the temperature change of these buffers within a 1-min lysis process is also expected to be very minor.

**Fig. 3.**
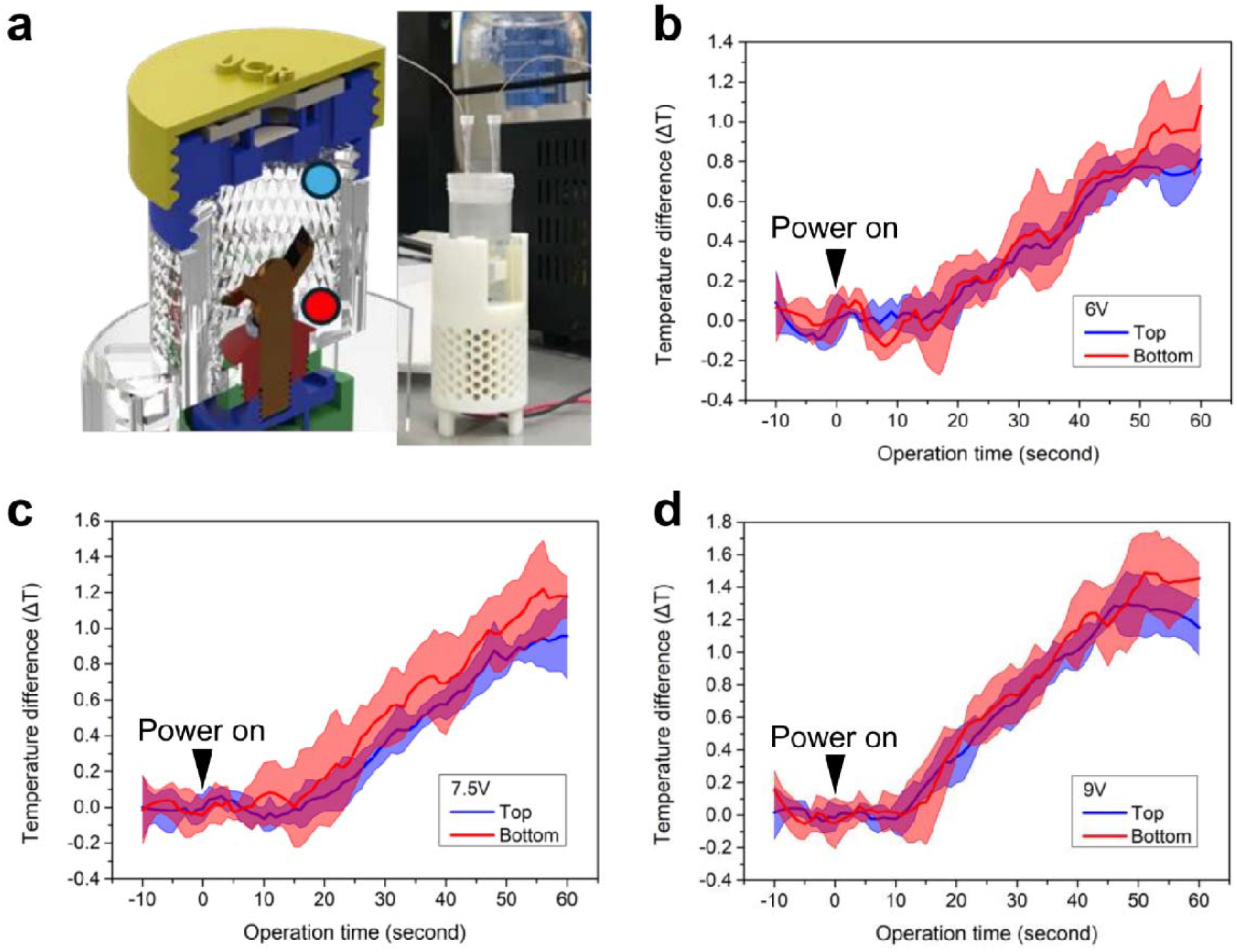
Evaluation of temperature inside lysis chamber during a 1-minute operation. (a) A cutaway view of the lysis chamber depicting the thermocouple locations where temperatures were measured and a photo showing the thermocouple wiring setup. The blue spot is above the blades (top), red is below (bottom). (b) Temperature profile at the top and bottom during 1-minute operation at 6V (N = 3). (c) Temperature profile at the top and bottom during 1-minute operation at 7.5V (N = 3). (d) Temperature profile at the top and bottom during 1-minute operation at 9V (N = 3).

#### 3.1.2 Evaluation of resin release

Since the entire handheld device is assembled with multiple 3D-printed parts made of resin, it was a concern that cured or uncured resins could be released into the lysis buffer during operation, affecting downstream extraction and detection. To evaluate this possibility, pristine guanidine lysis buffer was loaded in the chamber and the motor was run for 1 minute at 6, 7.5, and 9V, respectively. The buffers were then collected and analyzed using attenuated total reflectance Fourier transform infrared spectroscopy (ATR-FTIR) to characterize signals from specific functional groups of the resins in the buffer. Additionally, two further conditions were included to rule out any other signals due to heating the lysis buffer and the use of the sealing silicone grease.

Based on the data sheets provided by the manufacturer, Veroclear-RGD810 and VeroWhitePlus-RGD835 mainly contain exo-1,7,7-trimethylbicyclo[2.2.1]hept-2-yl acrylate and tricyclo decane dimethanol diacrylate, where acrylic group dominates (Stratasys Ltd. 2016a, b). As previously reported, the acrylic group is expected to show peaks at around 1720 cm^-1^ (i.e., 1710, 1718, 1720, 1723 cm^-1^), indicating C=O stretching. These peaks are straightforward evidence of the carbonyl group in acrylate (Loos et al., 2021; Rimington et al., 2018; Wajahat et al., 2024; Wang et al., 2017), which is not observed in any of these cases (Fig. S4). Additionally, characteristic peaks contributed by various vibrations of the vinyl group, C=C (810, 900, 910, 980, 1180, 1400, 1615, 1620-1640, 1643, 2975, and 3080 cm^-1^), are also important indicators to determine the existence of acrylic resins (Breitmoser et al., 2022; Chen et al., 2017; Kim et al., 2017; Macdonald et al., 2017; Rimington et al., 2018; Rosa 2023). However, none of the peaks are particularly obvious in these samples either, as shown in highlighted region i. Peaks pointing downward at 2855 cm^-1^ and 2920 cm^-1^ shown in the spectra might be inferred as the reduction of functional groups with =C−H stretching (Loos et al., 2021; Rimington et al., 2018). Since the absorbances are relatively small and remain unchanged across different samples, it could be a background subtraction issue due to existing molecules being interfered with by the water broadband next to these peaks, as shown in highlighted region ii (2900-3700 cm^-1^, O−H stretching) (Higl et al., 2021).

In fact, peaks in the spectra are mostly in the range 1950-2170 cm^-1^, with a major one at 1650 cm^-1^. Notably, it is observed that the changes of absorbance at 1650 cm^-1^ and 2060-2070 cm^-1^ are synchronous and seem inversely proportional to motor voltages. It is possible that guanidinium thiocyanate and PVP molecules may adsorb to the surface of the resin chamber, leading to the decrease of the absorbances at 1650 cm^-1^ (C=O stretching of PVP and C=N stretching of guanidinium thiocyanate) and 2060-2070 cm^-1^ (S−CN stretching of guanidinium thiocyanate) (Bramanti et al., 2022; Heyda et al., 2017; Zucchiatti et al., 2018). The rest of the peaks within the range (1950-2170 cm^-1^) might be correlated with asymmetric stretching of N=C=S (Conrad et al., 2021; Kyriakou et al., 2022). This analysis suggests that resin is not released into the lysis buffer during operation. The possible adsorption of the guanidinium thiocyanate to the resin, however, did not seem to adversely impact the device’s performance.

#### 3.1.3 Optimization of the operation using the handheld device

To maximize the nucleic acid release of the device, two different lysis chambers were evaluated: one with smooth walls and one with knurled walls. Three healthy citrus species (sour orange, Limoneira, and Washington navel) were macerated at three different voltages (6, 7.5, and 9V) for 1 minute. RNA was then extracted using a MagMAX kit and subsequently analyzed in terms of quantification cycles (Cq) of the internal control, COX, and total DNA and RNA concentrations. Additionally, the quantitative results were compared side by side with those from samples prepared with a mortar and pestle.

As shown in Fig. 4a, sample lysis using the knurled chamber always resulted in higher RNA concentration than the smooth chamber, regardless of operation voltages, with up to 1.76 times higher concentrations at 7.5 V. While comparing these with their mortar and pestle (MP) controls, the use of a knurled chamber achieved RNA yields of up to 94.6% (9 V) and 78.4% (7.5 V) for Limoneira and Washington navel, respectively. In contrast, the highest achievable total RNA concentration using a smooth chamber was only 53.2% (9 V) of its MP control. The results indicate that a knurled chamber enables much higher lysis efficiency than a smooth chamber, even when operated at a lower voltage. The amount of nucleic acids extracted using a knurled chamber was found to be quite consistent across different citrus species (Fig. 4a).

**Fig. 4.**
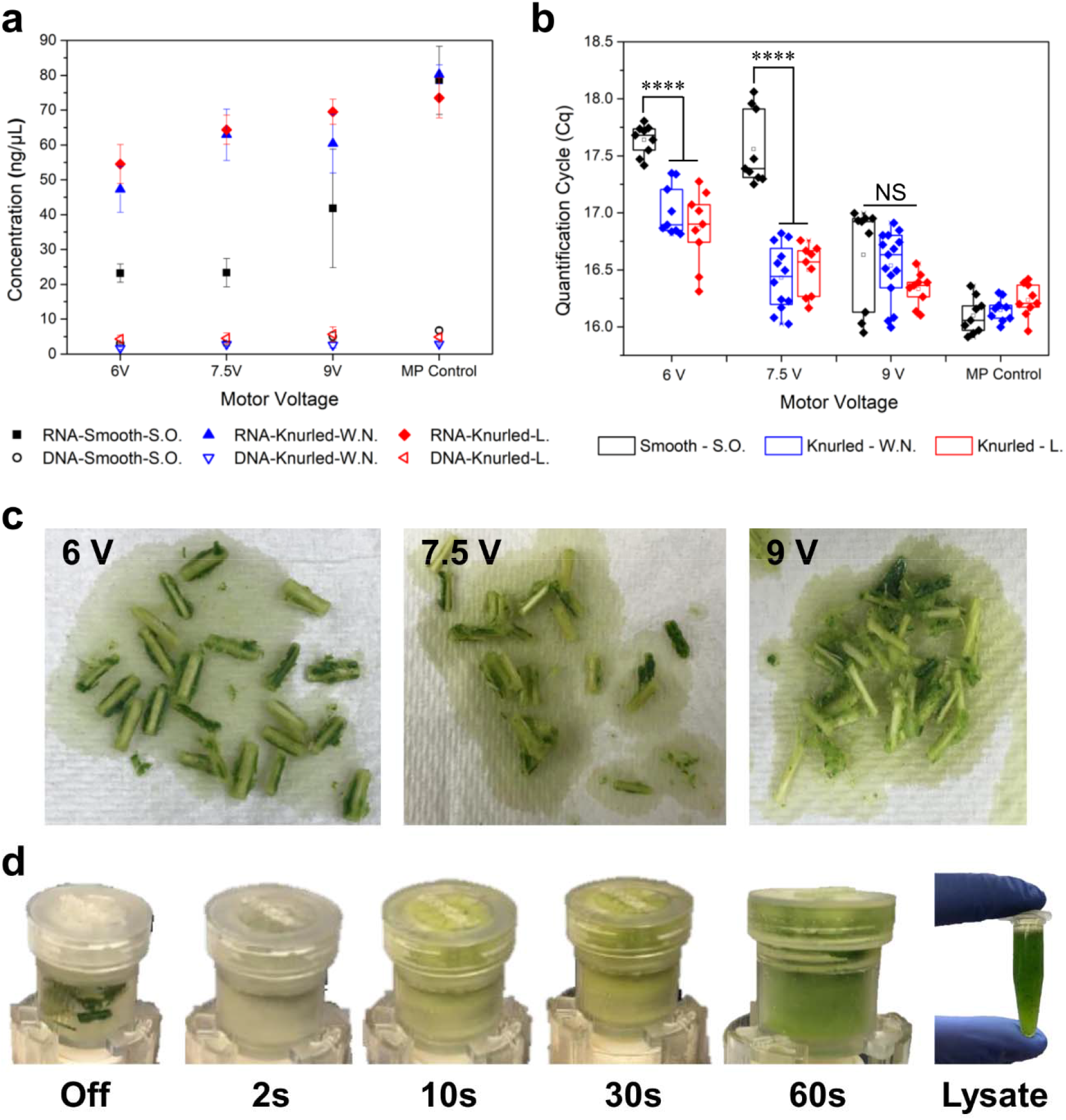
Optimization of nucleic acid release in terms of motor voltages and chamber features. (a) Comparison of total RNA and DNA concentrations among different voltages and the mortar and pestle grinding benchmark (N = 4 for Washington navel, and N = 3 for the rest). (b) Comparison of Cq values among different operation voltages and the mortar and pestle grinding benchmark (N = 12 for Washington navel and N = 9 for the rest). (c) Sample tissues after 1-minute lysis. (d) Color change of the lysate during the 1-minute lysis process at 7.5 V. **** denotes p ≤ 0.0001 and NS (Not significant) denotes p > 0.05.

When comparing Cq values (Fig. 4b), the advantage of the knurled chamber is even more apparent, especially at 6 and 7.5 V. It is inferred that knurling enhances the pulverization of tissues while the blade is spinning at high speed. The effect of this pulverization after 1 minute of operation can be seen in Fig. 4c. However, at 9V, the performances of the knurled and smooth chambers are much more similar, likely due to the much higher speed of the blades at that voltage. Notably, no significant differences were found in the Cq values of Limoneira and Washington navel at 7.5 and 9 V. This suggests that operation at 7.5 V is an optimized condition, outputting equivalent yield with a lower power requirement. Visualization of the sample lysis at 7.5 V can be seen in Fig. 4d, where the lysate turned green within 10 seconds of operation. Based on the optimization outcome, a knurled chamber with 7.5 V was selected for use throughout th following study.

### 3.2 Evaluation of device reusability

To prevent cross-contamination between uses of the handheld device, it is critically important to eliminate any previous genetic material in the chamber. Thus, a sterilization protocol from a previous study was adopted and modified in terms of soaking time (Dang et al., 2022). Specifically, the handheld device was disassembled and washed with water to remove most solid residues and residual guanidine lysis buffer before soaking the device in a 10% household bleach solution (approximately 0.825% sodium hypochlorite). As shown in Fig. 5a, five different time lengths (15, 30, 45, 60, and 75 minutes) of bleach soaking were tested for the complete elimination of COX after each 1-minute maceration of healthy tissues. COX was selected as a proof-of-concept model due to its abundance in plant lysate. This was followed by a 5-minute soak in a 2.3% sodium ascorbate solution to neutralize the residual bleach and then a brief rinse in water before it was ready for the next sample (United States Department of Agriculture Forest Service 2005).

**Fig. 5.**
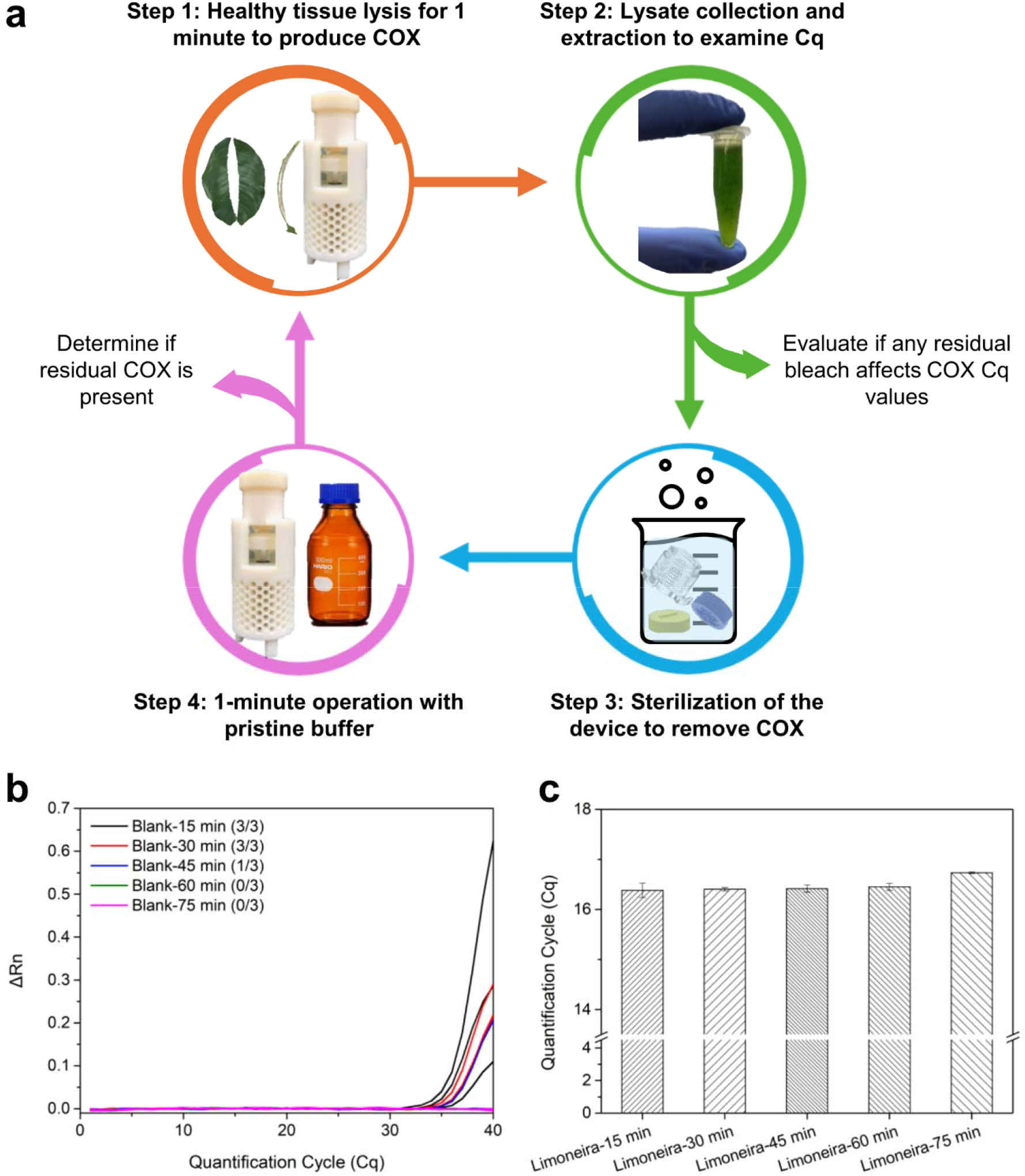
Evaluation of the sterilization protocol. (a) Stepwise procedure of the sterilization protocol. (b) RT-qPCR real-time curves of blank lysis extracts subjected to different bleach soaking times, where the number in the parathesis behind refers to a positive rate (N = 3). (c) COX Cq values of healthy extracts collected after different bleach soaking times (N = 3).

According to the results shown in Fig. 5b, it was observed that the COX gene was successfully eliminated after 60 minutes of soaking in the household bleach solution, whereas soaking between 15-45 minutes was insufficient for complete disinfection. Thus, a 60-minute soaking was selected as the optimized sterilization protocol and was subsequently tested with two different types of citrus pathogens to further assess its disinfection ability. Additionally, the possibility of residual bleach impacting the quality of released nucleic acids was assessed by comparing the COX Cq values of healthy lysates collected after each testing cycle. As shown in Fig. 5c, the quality of the COX gene was not adversely affected by residual bleach, as seen by the consistent Cq values (16.4 – 16.7) across the five samples.

Generally, this modified protocol effectively prevented cross-contamination and does not use ethanol, unlike the original protocol, where harmful byproducts could be produced if not carefully implemented (Dang et al., 2022). The use of sodium ascorbate for bleach neutralization is more eco-friendly and safer to handle than other dechlorination chemicals, which is especially suitable for disinfecting the used device in the field (United States Department of Agriculture Forest Service 2005).

### 3.3 In-device sample preparation for the detection of citrus pathogens

#### 3.3.1 Optimization of in-device sample preparation

To enable in-device sample preparation for downstream qPCR-based assays, the wash and elution protocols were modified from our previously reported paper-disk extraction protocol (Liu et al., 2024). Specifically, the number of wash steps and the volume of elution buffer were adjusted (Fig. 6a), whereas the wash/elution buffer composition stayed unchanged. Introducing a second wash step and doubling the elution buffer volume were both attempted to reduce the effects of residual RT-qPCR inhibitors. CTV was used as a proof-of-concept pathogen for these optimizations.

**Fig. 6.**
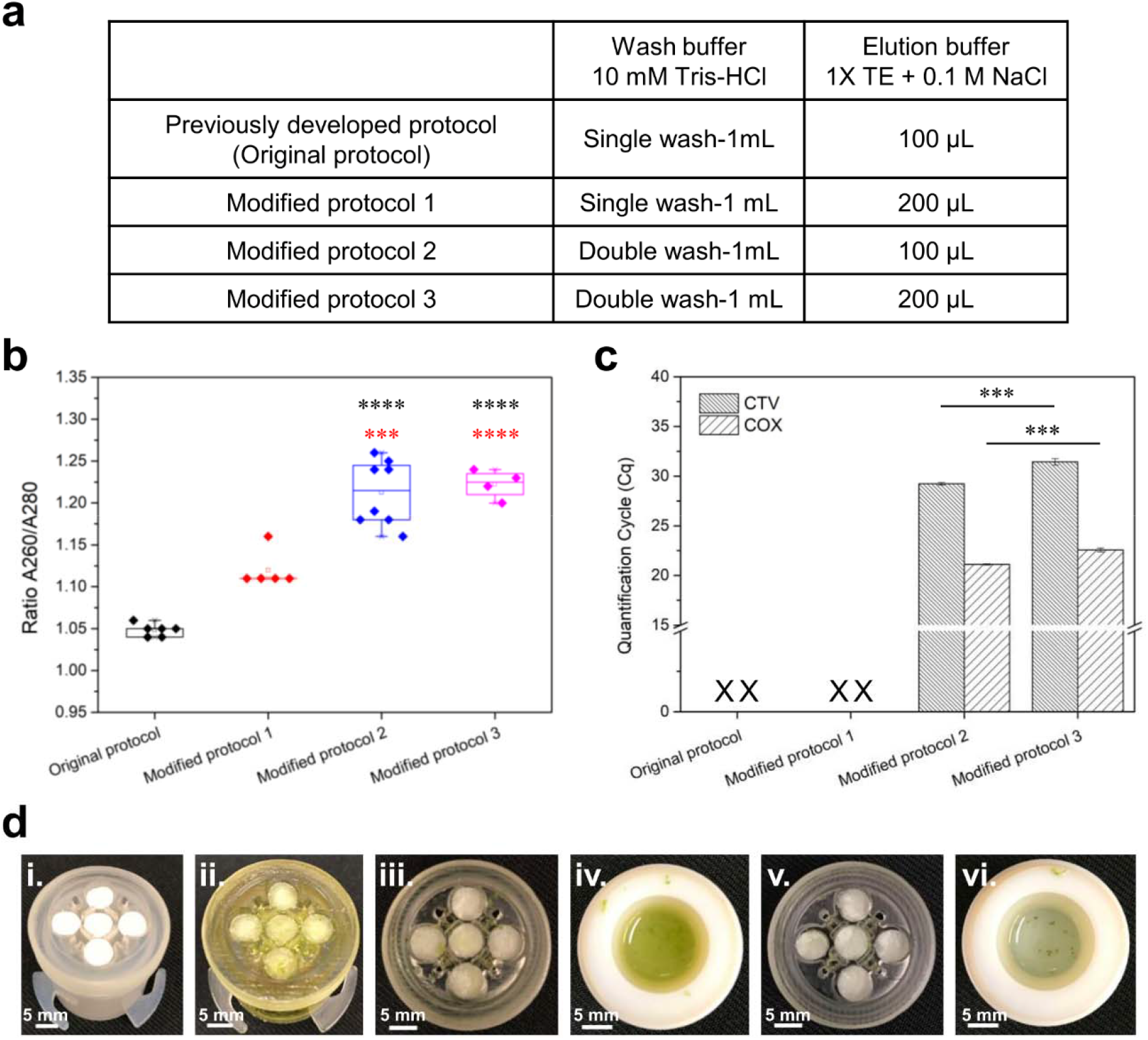
Optimization of the paper disk washing and elution protocol for in-device nucleic acid extraction. (a) The four tested paper disk washing and elution protocols. (b) Evaluation of the purity of different extracts in terms of A260/A280 ratio (N = 4 – 8). (c) Comparison of Cq value among the four tested protocols. Neither of the single-wash protocols resulted in a detectable signal (N = 3). (d) Paper disk washing sequence: i) before and ii) after soaking in crude lysate; iii) Paper disks after single wash; iv) Wash buffer after the first wash; v) Paper disks after double wash; vi) Wash buffer after the second wash. *** and **** denote p ≤ 0.001 and p ≤ 0.0001, respectively.

The purity of the resulting extracts was estimated by measuring the A260/A280 ratio of each extract, as shown in Fig. 6b. With a single-wash step, extracts from the original protocol and modified protocol 1 showed low purities, ranging from 1.04 – 1.16, compared to the MagMAX extract (2.09). Similarly, false-negative RT-qPCR results for CTV and COX were also observed (Fig. 6c). It is inferred that the RT-qPCR reactions were inhibited because contaminants, such as the lysis buffer or the plants’ proteins, remained in the paper disks and were not sufficiently diluted despite doubling the volume of elution buffer.

The extracts subjected to a double-wash step (i.e., modified protocols 2 and 3), both demonstrated higher A260/A280 ratios, ranging from 1.18 – 1.25, than the single-wash cases (Fig. 6b). Although the purities did not improve greatly and were not close to what is expected for a pure extract (1.8 – 2.0) (Thermo Scientific 2012); the positive RT-qPCR results of CTV and COX proved the double-wash protocol was sufficient in mitigating inhibition (Fig. 6c). Additionally, it was observed that doubling the volume of elution buffer increased Cq values of both CTV and COX (Fig. 6c). This suggests that further adjustments of elution volume should not be required as physically removing inhibitors via repeated washing steps is much more effective than diluting them for RT-qPCR assays. Ultimately, the double-wash protocol (protocol 2) was adopted for the subsequent tests for in-device sample preparation and detection of different citrus pathogens.

#### 3.3.2 Validation of the optimized in-device sample preparation with citrus pathogens

Using the above optimized protocols, the next step was assessing the handheld device’ applicability to different citrus pathogens. To achieve this goal, CTV and *S. citri* were selected as the representative models of two major categories, RNA and DNA pathogens. Particularly, MagMAX extracts directly prepared from the same pre-soak lysates served as a benchmark to evaluate the capability of in-device sample preparation in nucleic acid recovery.

To evaluate the amount of nucleic acids being recovered using in-device sample preparation, Cq differences and total RNA concentrations between the two extraction approaches were analyzed. In terms of the qPCR/RT-qPCR results (Fig. 7a-d), the average CTV Cq difference (N = 12) between MagMAX and in-device extracts was 7.34, whereas the average *S. citri* Cq difference (N = 12) was 2.40, which was much smaller than that of CTV. However, the average COX Cq differences of CTV (4.19) and *S. citri* (3.70) were close (Fig. 7a-d), indicating similar total nucleic acid recovery efficiency across two different diseased plants. It is inferred that *S. citri* (DNA) is more tolerant to the 24-hour air drying process than CTV (RNA), which is more vulnerable to ambient enzymes (e.g., RNase). Microscopic interaction among these extracted proteins and total nucleic acids, and paper substrate in terms of size, structure, and charge distribution could be another factor, leading to different preferences between *S. citri* and CTV in physically binding to paper fibers (Pelton 2009; Seok et al., 2019). Although there is not any false negative being observed within these paper-disk extracts for all the CTV and S. citri assays in this test, the Cq delay compared to benchtop extracts still implied that a further improvement in nucleic acid extraction is needed, especially when processing infected plants with low viral load.

**Fig. 7.**
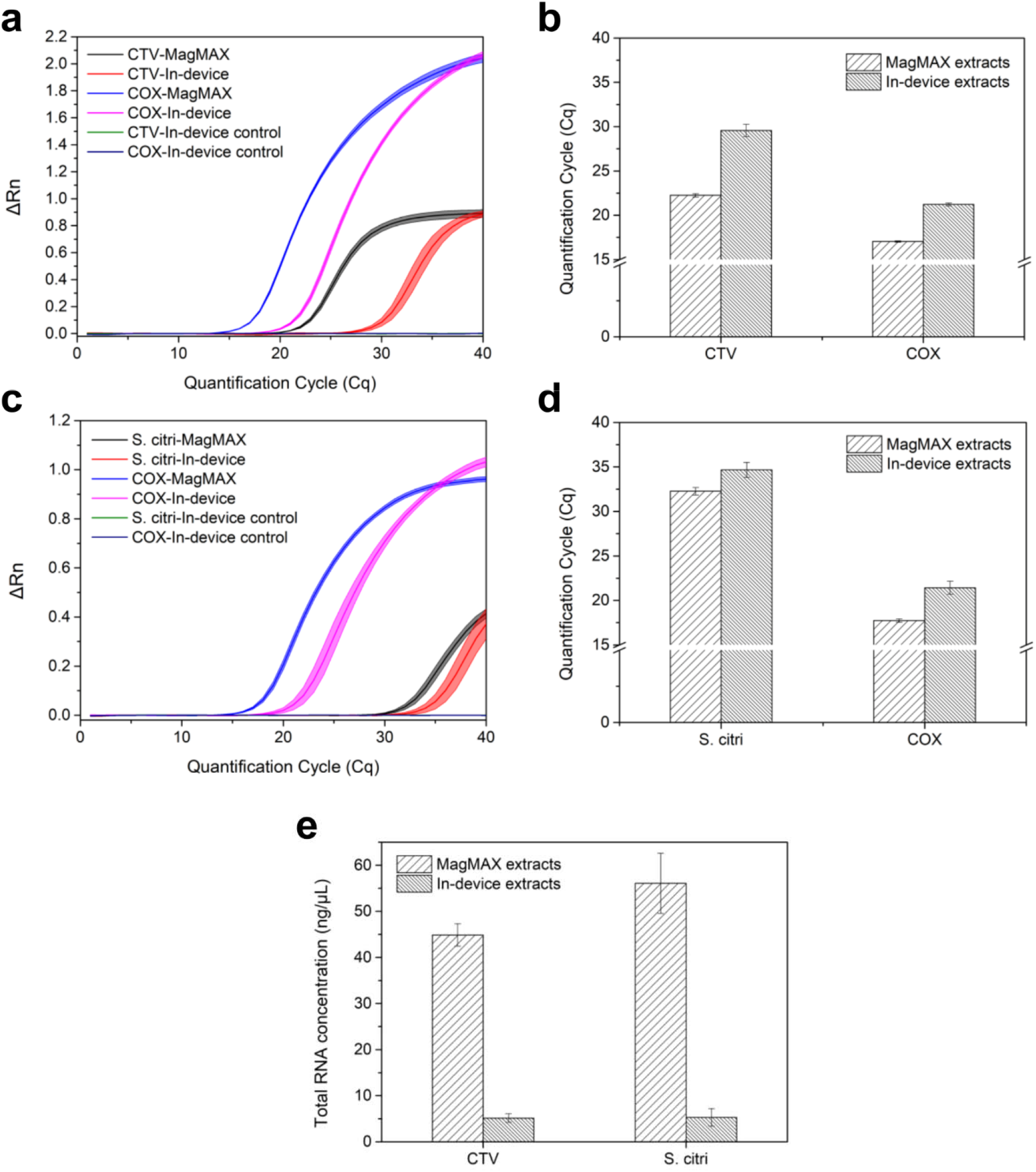
Validation of the optimized in-device sample preparation with citrus pathogens. (a) RT-qPCR curves of MagMAX and in-device extracts from CTV-infected source (N = 12). (b) Comparison of CTV and COX Cq values between MagMAX and in-device extracts (N = 12). (c) qPCR curves of MagMAX and in-device extracts from *S. citri*-infected source (N = 12). (d) Comparison of *S. citri* and COX Cq values between MagMAX and in-device extracts (N = 12). (e) Comparison of total RNA concentration between MagMAX and in-device CTV and *S. citri* extracts (N = 12).

It was observed that in-device extracts from both *S. citri* and CTV-infected plants yielded similar total RNA at around 5.18-5.29 ng/µL (Fig. 7e). These are approximately 10% of the total RNA yield of MagMAX extracts (Fig. 7e). Total DNA yields, however, were not reliably detectable using the spectrophotometer, but *S. citri* DNA was detectable via qPCR. This suggests that the capability of a single paper disk to retain nucleic acids is the limiting factor, rather than the device’s capability of releasing nucleic acids from lysed plant tissues.

Additionally, assay variations were evaluated further to confirm the reliability of the in-device sample preparation strategy. According to the qPCR/RT-qPCR results (Fig. 7b, 7d), both in-device CTV and *S. citri* sample preparation demonstrated low intra-assay (< 3.1%) and low inter-assay (< 3.8%) coefficients of variation (CV), indicating high consistency and reproducibility.

### 3.4 Conclusion

In this study, a 3D-printed handheld device was designed and fabricated for quick sample preparation of citrus leaves for pathogen detection. The operation of the 3D-printed handheld device was optimized in terms of motor voltage and chamber features, where using a knurled chamber and a 7.5 V motor voltage for just 1 minute of operation, yielded up to 87.6% of the total RNA extracted using a traditional mortar and pestle grinding protocol. Other aspects of the device were also investigated, such as heat generation during operation and the potential release of resin from the 3D-prints were found to have little-to-no impact on the quality of the extracted nucleic acids. Further, sterilization protocols were investigated, and it was determined that soaking the device in 10% household bleach solution for 60 minutes, followed by a brief wash with 2.3% sodium ascorbate solution and water, effectively eliminated all remaining pathogenic nucleic acids in the lysis chamber. The in-device sample preparation protocols were also optimized and verified with citrus sources infected with RNA and DNA citrus pathogens (i.e., CTV and *S. citri*) in qPCR-based assays, achieving low intra- and inter-assay variations (3.1% and 3.8%, respectively). The results prove that 1-minute mechanical maceration using such a small handheld device can perform almost equivalently in sample lysis to manual grinding using a mortar and pestle. Particularly, in-device sample preparation integrated with the paper disks showed good reliability and compatibility across different pathogens and citrus varieties, suggesting its potential to be adopted to process more plant for disease diagnoses.

This portable, easy-to-use device is a potential alternative to standard lab protocol, allowing individuals to rapidly prepare ready-to-use nucleic acids in the field for analysis. Current disease diagnosis practices can be expedited without the need for cumbersome specimen collection, packaging, and submission. Particularly, it shortens the turnaround time required from sample to answer, where the only end products, dried paper disks, can either be shipped directly to the lab for further analysis or be eluted locally for on-site detection. Although untreated chromatography paper was sufficient for the current study, using different materials or other chemically modified membranes with better nucleic acid recovery may be preferred when dealing with non-symptomatic plants for early-stage diagnosis (Tang et al., 2022). This innovative device significantly improves sample lysis efficiency and streamlines nucleic acid preparation, providing a reliable and expedient alternative for in-field molecular biology applications.

## CRediT authorship contribution statement

**Chia-Wei Liu:** Conceptualization, Methodology, Validation, Formal analysis, Investigation, Writing – original draft, Writing – Review & Editing, Visualization. **Brent Kalish:** Conceptualization, Methodology, Writing – original draft, Writing – Review & Editing, Visualization. **Sohrab Bodaghi:** Conceptualization, Methodology, Resources, Writing – Review & Editing, Supervision, Funding acquisition. **Georgios Vidalakis:** Conceptualization, Resources, Writing – Review & Editing, Supervision, Project administration, Funding acquisition. **Hideaki Tsutsui:** Conceptualization, Writing – Review & Editing, Supervision, Project administration, Funding acquisition.

## Acknowledgment

The authors thank Dr. Jun Sheng’s research group and Boshen Qi for providing access to their 3D printers. The authors thank Dr. Arunabha Mitra for providing access to his citrus plants in Agricultrual Operations (AgOPs) at university of California, Riverside. The authors thank Dr. Lingchao Zhu at UCR Analytical Chemistry Instrumentation Facility (ACIF) for his assistance in ATR-FTIR measurements.

## Data Availability

The data supporting the findings of this study are available from the corresponding author upon reasonable request.

