## Supplementary Data for "A 3D-printed handheld device for quick citrus tissue lysis and nucleic acid extraction"

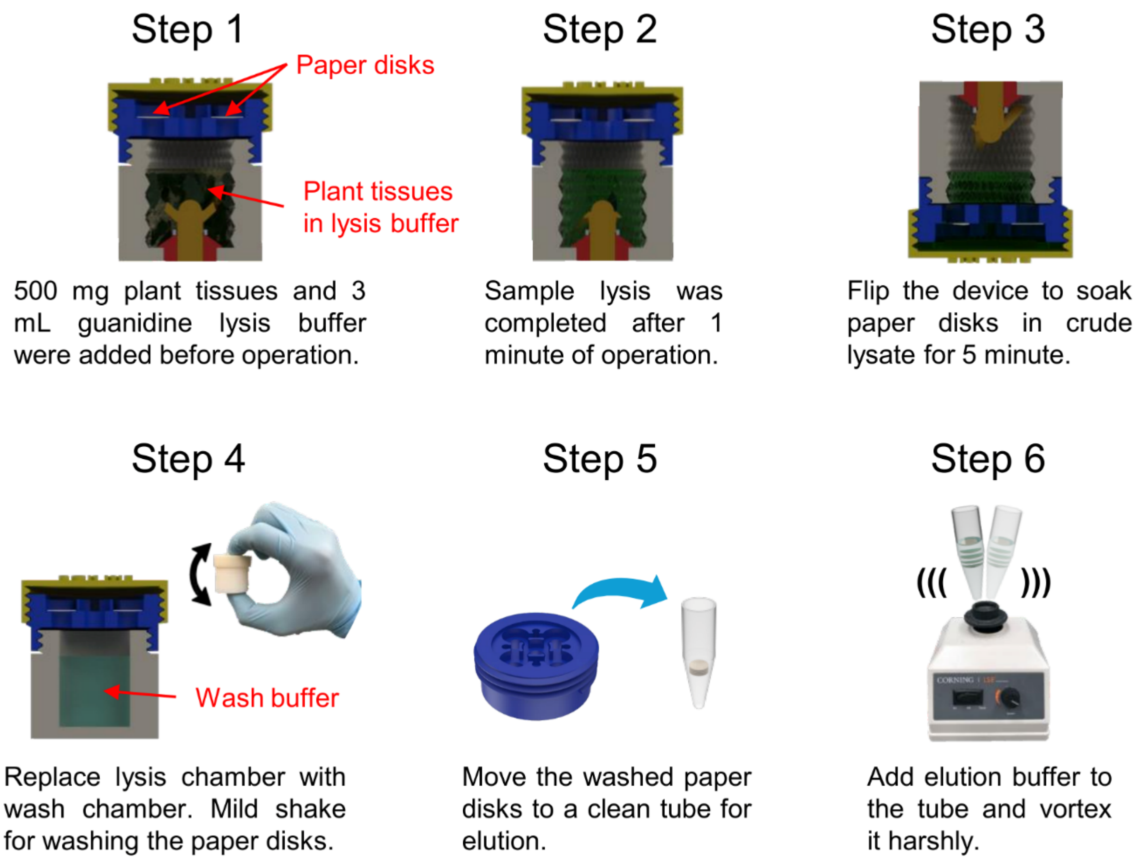


Fig. S1. Stepwise procedure for sample preparation using the handheld device.

**a**

| | 6 V | | | 7.5 V | | | 9 V | | |
|---|---|---|---|---|---|---|---|---|---|
| Before operation (3 mL lysis buffer added) | 3.4 g | | | 3.4 g | | | 3.4 g | | |
| 1-min operation w/o silicone grease | 3.19 | 3.03 | 3.02 | 2.36 | 2.35 | 2.42 | 2.26 | 2.57 | 2.73 |
| 1-min operation w/ silicone grease | 3.24 | 3.25 | 3.33 | 3.28 | 3.05 | 3.30 | 2.90 | 3.19 | 3.10 |

**b**

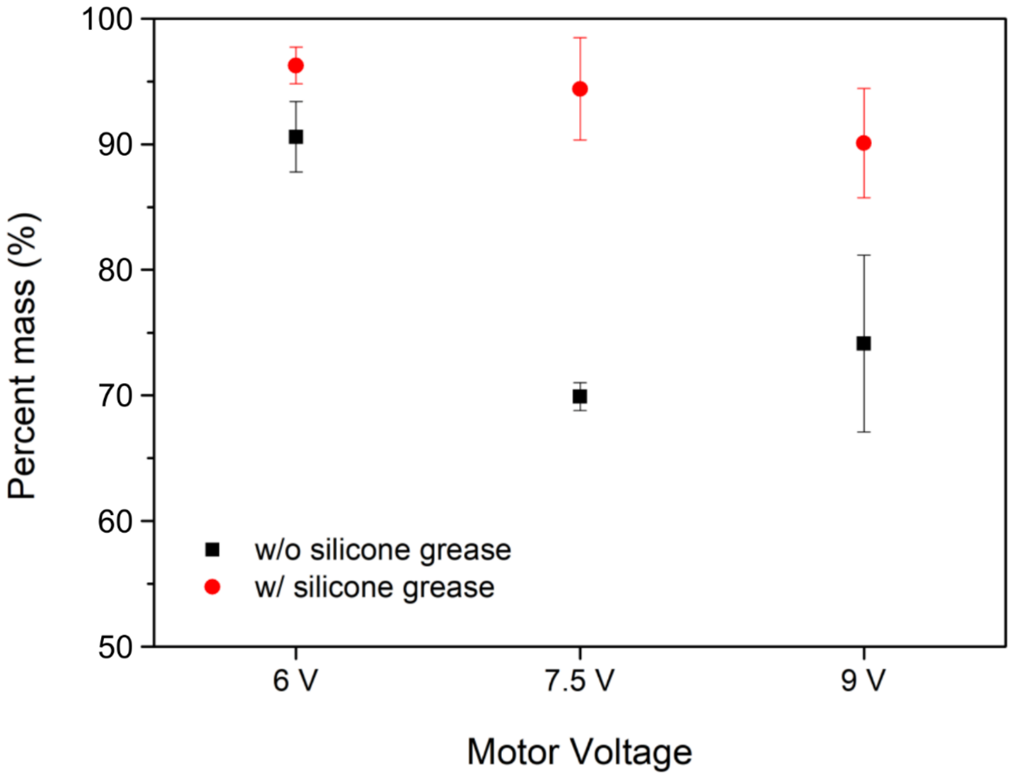


Fig. S2. Buffer leakage with and without using silicone grease around the shaft of the blade (N = 3). (a) The mass of lysis buffer before and after each test. (b) Percent mass loss of lysis buffer before and after application of the silicone grease under different motor voltages.

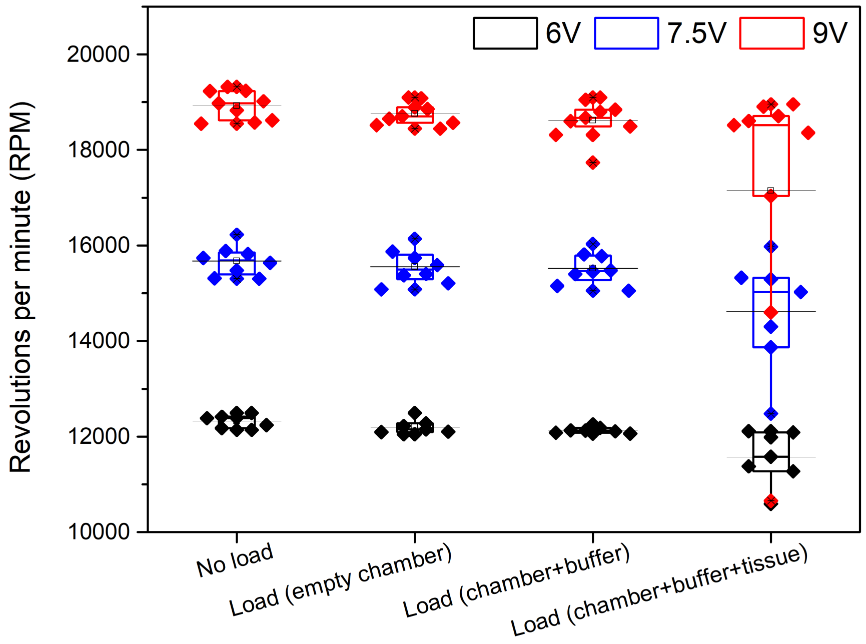


Fig. S3. The change in RPM under different loading conditions and motor voltages. The rotational speeds generated by the different voltages were consistent across the different loading conditions until the introduction of plant tissues.

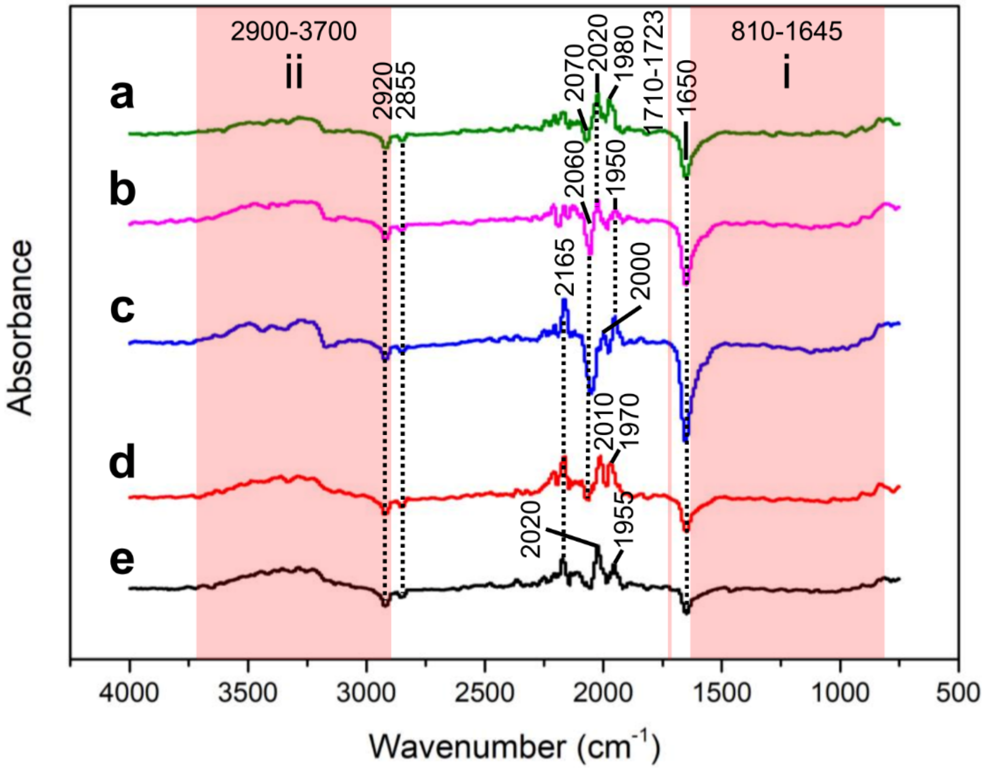


Fig. S4. Comparison of ATR-FTIR spectra evaluating the possible release of resin molecules and other chemicals. Band i) shows no distinct peaks that could be correlated with possible resin indicators, while band ii) corresponds to the water broadband which masks any potential minor peaks. (a) Guanidine lysis buffer subject to 1-minute operation at 9 V. (b) Guanidine lysis buffer subject to 1-minute operation at 7.5 V. (c) Guanidine lysis buffer subject to 1-minute operation at 6 V. (d) Heating control: Guanidine lysis buffer subject to 1 minute of heating at 30°C without motor operation. (e) Silicone grease control: Guanidine lysis buffer mixed with silicone grease without motor operation.

Table S1. Non-inoculated and inoculated source plants used in the study

| Category | Source plant | Pathogens |
|---|---|---|
| Healthy | Sour orange | Non-infected |
| | Washington navel | |
| | Limoneira | |
| Single-infected | Pineapple sweet orange | *Citrus tristeza virus* (RNA virus) |
| | Sweet orange | *Spiroplasma citri* (DNA bacterium) |

Table S2. Oligonucleotide primers and probes used for the qPCR/RT-qPCR assays in this study.

| Isolates | Primers/probes | Sequence 5' – 3' | Nucleotide position | Amplicon size (bp) | Ref. |
|---|---|---|---|---|---|
| COX | Forward | AATCTGACCTTCT TTCCCATGC | 32-53 | 162 | [1, 2] |
| | Reverse | AAGTGATTGTTAC GACCACGAAGA | 194-171 | | |
| | Probe | ATCCAGATGCTTA CGCTGG | 96-114 | | |
| CTV | Forward | TGTGTGCGGATTT CTTGACTG | 701-721 | 135 | [2] |
| | Reverse | TTCCCAAGCTGCC TGACATT | 833-814 | | |
| | Probe | AAGCGAGGGGCT GAT | 784-798 | | |
| *S. citri* | Forward 1 | ATTGCAGCACCTG CAACTGTAG | 112-133 | 114 | [3] |
| | Reverse | TGTTTTTACAACT CCTTGCACTGC | 225-202 | | |
| | Probe | ACAGCGTTAGAA GCTAAT | 175-192 | | |

Nucleotide position is based on GenBank accessions: COX-CX297817, CTV-M76485, and S. citri-FJ755921.

---------------------------------------------------------------------------------------------------------------------------------------

[1] Osman, Fatima, et al. *Journal of Virological Methods* 220 (2017): 40-52.

[2] Osman, Fatima, et al. *Journal of Virological Methods* 245 (2015): 64-75.

[3] Shi, Jinxia, et al. *Phytopathology* 104.2 (2014): 188-195.

Table S3. Kits and protocols used for the qPCR/RT-qPCR assays in this study.

| Pathogen | Assay | Composition | Thermal-cycling protocol |
|---|---|---|---|
| *S. citri* | Singleplex qPCR iTaq™ Universal Probes One-Step Kit (Bio-Rad, USA) | 6 µL 2X PCR buffer<br>0.576 µL primer/probe mix (300 nM for forward primer, 600 nM for reverse primer and 200 nM for probe as final concentrations)<br>3.424 µL ddH2O<br>2 µL DNA template. | Incubation stage (2 min at 50 °C and 10 min at 95 °C) and 40 repeating cycles (15 sec at 95 °C and 1 min at 60 °C). |
| CTV | Singleplex RT-qPCR AgPath-ID One-Step RT-PCR kit (Thermo Fisher Scientific, USA) | 6.25 µL 2X RT-PCR buffer<br>0.6 µL primer/probe mix (417 nM for primers and 83 nM for probe as final concentrations)<br>0.5 µL 25X RT-PCR enzyme mix<br>2.65 µL ddH2O<br>2 µL RNA template. | Incubation stage (10 min at 45 °C and 10 min at 95 °C) and 40 repeating cycles (15 sec at 95 °C and 45 sec at 60 °C). |
